# A tractable *Drosophila* cell system enables rapid identification of *Acinetobacter baumannii* host factors

**DOI:** 10.1101/2020.04.09.034157

**Authors:** Qing-Ming Qin, Jianwu Pei, Gabriel Gomez, Allison Rice-Ficht, Thomas A. Ficht, Paul de Figueiredo

## Abstract

*Acinetobacter baumannii* is an important causative agent of nosocomial infections worldwide. The pathogen also readily acquires resistance to antibiotics, and pan-resistant strains have been reported. *A. baumannii* is widely regarded as an extracellular bacterial pathogen. However, accumulating evidence demonstrates that the pathogen can invade, survive or persist in infected mammalian cells. Unfortunately, the molecular mechanisms controlling these processes remain poorly understood. Here, we show that *Drosophila* S2 cells provide several attractive advantages as a model system for investigating the intracellular lifestyle of the pathogen, including susceptibility to bacterial intracellular replication and limited infection-induced host cell death. We also show that the *Drosophila* system can be used to rapidly identify host factors, including MAP kinase proteins, which confer susceptibility to intracellular parasitism. Finally, analysis of the *Drosophila* system suggested that host proteins that regulate organelle biogenesis and membrane trafficking contribute to regulating the intracellular lifestyle of the pathogen. Taken together, these findings establish a novel model system for elucidating interactions between *A. baumannii* and host cells, define new factors that regulate bacterial invasion or intracellular persistence, and identify subcellular compartments in host cells that interact with the pathogen.

## Introduction

*Acinetobacter baumannii* is a clinically important pathogen that can survive on hospital equipment and can cause serious nosocomial infections. The organism also causes serious wound infections in injured combat soldiers and in victims of traumatic injury. In addition, the bacterium can readily acquire multidrug, extensive-drug and even pan-drug resistance phenotypes. These attributes render *A. baumannii* a potential global threat to health-care settings (Harding et al., 2018; Mortensen and Skaar, 2013). *A. baumannii* is widely regarded as an extracellular bacterial pathogen; however, accumulating evidence indicates that the pathogen can invade and persist within an iron-starved compartment of mammalian cells (Harding et al., 2018; Mortensen and Skaar, 2013). In the past two decades, progress has been made in identifying and characterizing host factors that regulate the intracellular lifestyle of diverse pathogens, including *A. baumannii* (An et al., 2019; Choi et al., 2008; Parra-Millan et al., 2018; Rumbo et al., 2014; Smani et al., 2012; Wang et al., 2016). However, this aspect of the infection process remains poorly understood.

The evolutionarily divergent *Drosophila* S2 cell model system has been exploited as an alternative host system for studying mammalian host-pathogen interactions since it recapitulates conserved aspects of innate immunity (Criscitiello and de Figueiredo, 2013; Kim and Kim, 2005; Pandey et al., 2014). We previously demonstrated that mammalian orthologs of hits identified in RNAi (RNA interference) screens of the *Drosophila* S2 cells for host factors mediating pathogen infection are important for bacterial and fungal infection of mammalian cells, thereby validating the utility and convenience of this insect cell model for host-pathogen interaction studies (Pandey et al., 2018; Qin et al., 2011; Qin et al., 2008). The combination of the *Drosophila* S2 cell system and RNAi technology for depletion of host gene expression has also provided unprecedented opportunities for rapid functional elucidation of host factors. Here, we show that *Drosophila* S2 cells provide a convenient system for interrogating interactions between *A. baumannii* and host cells. We demonstrate the utility of this system by showing its use for identifying a role for host MAP kinase proteins in conferring susceptibility to intracellular parasitism. Ultimately, these findings may facilitate the development of novel host-directed anti-infectives for combatting the bacterium.

## Results

### *A. baumannii* infection induces host cell death in alveolar macrophages but not lung epithelial cells

The lung is an important site of *A. baumannii* infection. We therefore used gentamicin protection assays to determine whether alveolar macrophage (MH-S) or lung epithelial (TC-1 or MLE-12) cells were susceptible to *A. baumannii* intracellular parasitism. We found that the pathogen efficiently invaded alveolar cells (**Fig. 1A**). To confirm our findings, we used gentamicin protection assays to analyze intracellular persistence or replication of the pathogen during a time course of infection. CFU analysis demonstrated the replication and/or persistence of the bacteria in alveolar cells (**Fig. 1B-D**). Previous studies showed that *A. baumannii* invasion of mammalian cells can ultimately induce host cell death (Choi et al., 2005; Kim et al., 2008; Smani et al., 2011; Tamang et al., 2011). To determine whether the bacterium similarly killed alveolar cells, we measured the release of LDH, an indicator of host cell death, following infection (Pei and Ficht, 2004; Pei et al., 2008). We found that invasive bacteria did indeed induce the death of alveolar macrophages at 48 hr post infection (h.p.i.) in an MOI (multiplicity of infection)-dependent fashion (**Fig. 1D; Fig. S1**). These data show that *A. baumannii* invades, persists, or replicates in, and ultimately kills, alveolar macrophage cells.

**Figure 1.**
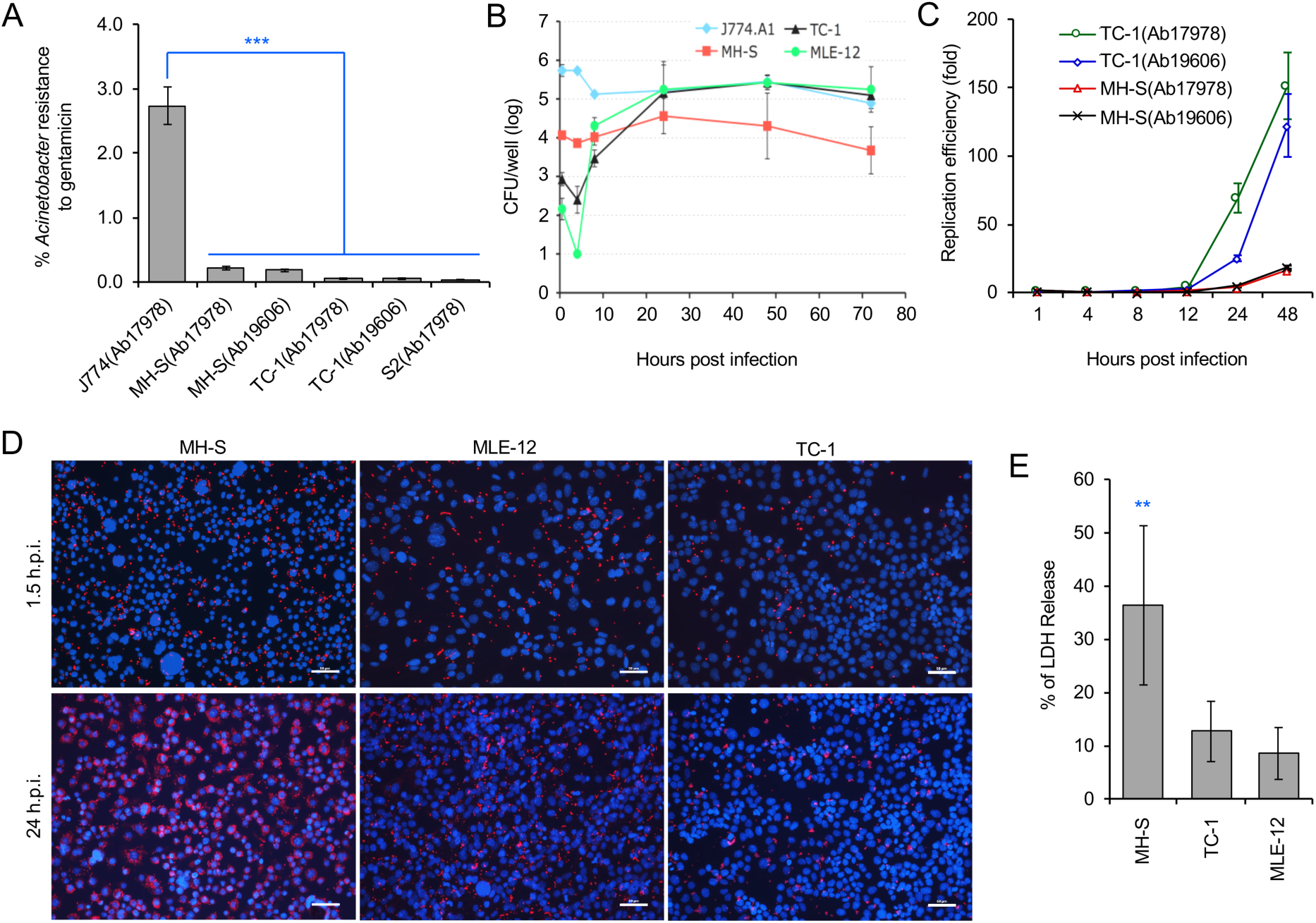
*Acinetobacter baumannii* invades and persists or replicates within mouse alveolar cells during a time course (72 hr) of infections. (A) Successful invasion of A. *baumannii* into various host cells. S2: *Drosophila melanogaster* S2 cells; Ab17978, Ab19606: A. *baumannii* wild-type strains 19606 and 17978, respectively. (B) Dynamics of intracellular Ab19606 replication or persistence in the indicated mouse cell lines. (C) Intracellular replication efficiency of *A. baumannii* in mouse alveolar epithelial (TC-1) or macrophage (MH-S) cells. (D) Representative images showing intracellular replication or persistence of *A. baumannii* in the indicated mammalian cells at 24 h.p.i.. (E) *A. baumannii* infection induces varying levels of cell death in alveolar cells (MH-S, TC-1 or MLE-12) at 48 h.p.i. Data represent means ± standard deviation (SD) from three independent experiments with triplicate wells examined for each treatment

### *A. baumannii* infects *Drosophila* S2 cells

*A. baumannii* enters membrane bounded vacuoles of epithelial cells in a process that depends upon the activities of microfilaments, microtubules, clathrin and β-arrestins (Choi et al., 2008; Smani et al., 2012). We sought to identify additional host mechanisms that support the intracellular survival or persistence of *A. baumannii*. However, bacterial invasion of mammalian cells induces cell death (**Fig. 1D; Fig. S1**) (Choi et al., 2005; Choi et al., 2008; Smani et al., 2012), an outcome which had the potential of confounding efforts to identify novel host mechanisms. We therefore tested whether *Drosophila* S2 cells, a macrophage-like cell line and a model system previously shown to be useful for elucidating host-pathogen interactions (Cherry, 2008; Pandey et al., 2014; Qin et al., 2011; Qin et al., 2008), displayed resistance to bacteria-induced killing. We found that *A. baumannii* efficiently invaded S2 cells, which were also resistant to pathogen-induced killing. Moreover, S2 cells were permissive to intracellular replication of *A. baumannii* (**Fig. A1; Fig. S2**). Similar to previous findings from mammalian host cell systems (Parra-Millan et al., 2018), *A. baumanni* resided in lysosomal marker LAMP-1 or cathepsin D-positive compartments at later time points post-infection (24 h.p.i.) (**Fig. S3**). Interestingly, intracellular bacteria were also located in ER (endoplasmic reticulum) marker calreticulin-positive vacuoles (**Fig. S3**). These data demonstrated that *Drosophila* S2 cells support *A. baumannii* infection. We therefore performed several experiments to determine the suitability of *Drosophila* S2 cells for identifying host factors that control the intracellular survival or persistence of *A. baumannii*.

### *Drosophila* S2 cells can be used as an alternative host to rapidly identify host factors that control the invasion, intracellular survival or persistence of *A. baumannii*

To identify host functions that confer susceptibility to *A. baumannii* internalization, survival or intracellular replication, we pre-treated S2 cells with compounds that disrupt host cell functions, and then measured levels of bacterial internalization or intracellular replication. We found that pre-treatment with brefeldin A (BFA), a molecule that disrupts ER to Golgi anterograde transport and the morphology and function of the trans-Golgi network and endosomes (de Figueiredo et al., 1998; de Figueiredo et al., 2000), altered the internalization, but not the replication, of the pathogen (**Fig. 2A, B**). Pre-treatment of host cells with myriocin, which depletes host cells of sphingolipids, did not alter either of these processes (**Fig. 2A, B**). However, host cells pre-treated with cytochalasin D, a compound that depolymerizes host cell actin, or wortmannin (WM), a class III phosphatidylinositol 3-kinase (PI3K) inhibitor, or SP600125, an inhibitor of the MAP kinase JNK, limited the internalization and replication of the pathogen (**Fig. 2A-B**). Interestingly, host cells pre-treated with bafilomycin A (BAF), a commonly used compound that inhibits autophagy by targeting lysosomes, limited the internalization but increased intracellular replication efficiency of the pathogen (**Fig. 2A-B**; **Fig. S4**). Importantly, these compounds did not disturb bacterial grow or cause cytotoxicity on host cells (Pandey et al., 2018; Qin et al., 2011; Qin et al., 2008), thereby suggesting that the observed reductions in pathogen internalization or intracellular replication did not emerge as a consequence of the drug acting directly upon the pathogen or host cells. Taken together, the data indicate that autophagosome biogenesis and accumulatio, as well as JNK activity, conferred susceptibility to *A. baumannii* infection of *Drosophila* S2 cells.

**Figure 2.**
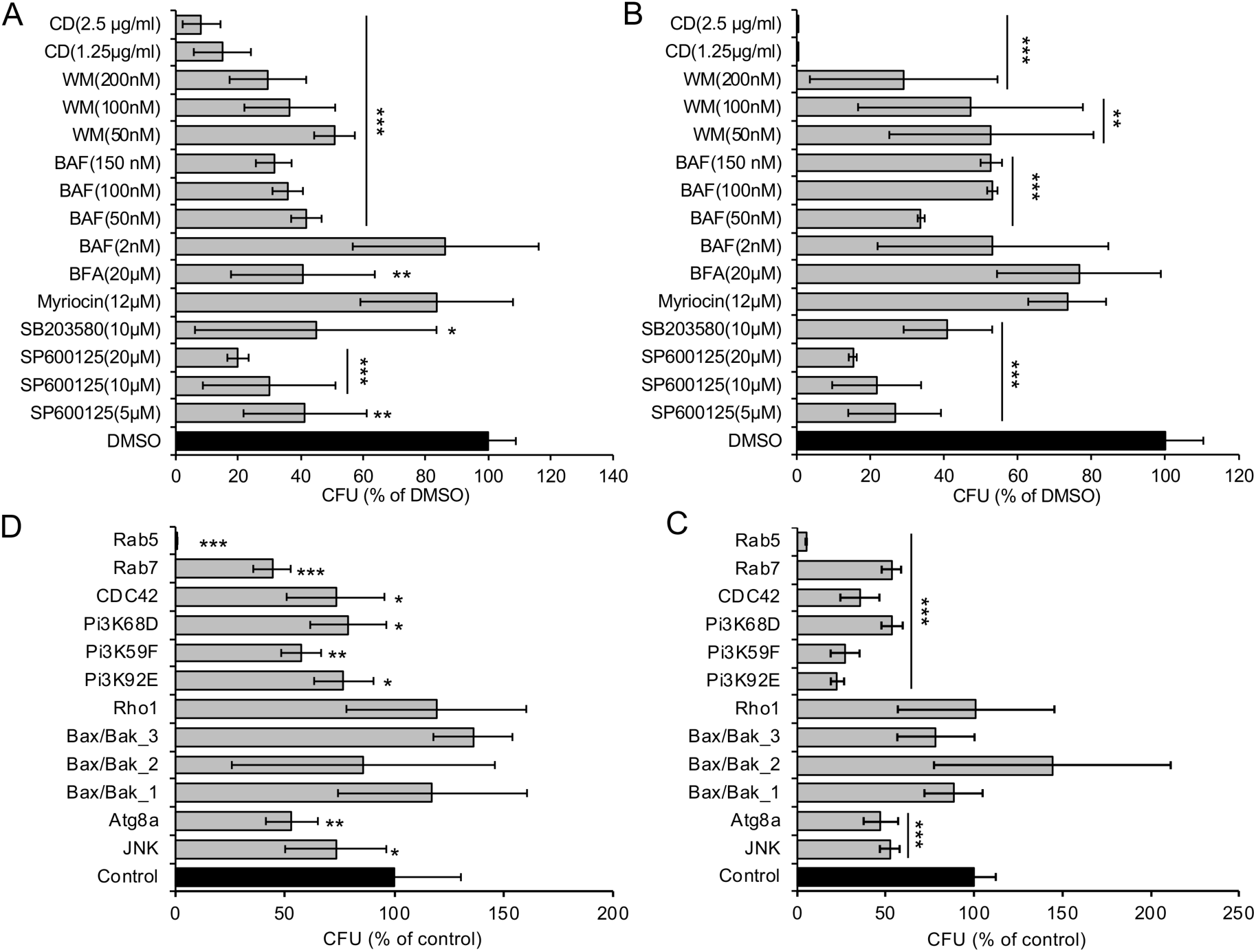
*A. baumannii* invasion and intracellular replication in *D. melanogaster* S2 cells pretreated with pharmacological compounds or treated with dsRNAs to deplete host proteins. (A, B) Drug-treated or untreated S2 cells were infected with Ab17978. At 1.5 (A) or 48 (B) h.p.i., the infected host cells were lysed for CFU analysis. (C, D) S2 cells were depleted of the indicated target genes using RNAi approaches and then infected with Ab17978 for 1.5 (C) or 48 (D) hr. Samples were then lysed for CFU analysis. Data represent means ± SD from four independent experiments with triplicate wells examined for each treatment. *, **, ***: significance at p < 0.05, 0.01 and 0.001, respectively.

To confirm and extend findings uncovered in our pharmacological analysis, we used RNAi technology to knockdown the expression of selected host target genes. S2 cells that had been treated with dsRNAs that targeted Rab5 and Rab7, which control the biogenesis of early and late endosomes, respectively, displayed resistance to the internalization and intracellular replication or persistence of the pathogen (**Fig. 2C-D**). Similarly, depletion of proteins encoding components of the PI3K complex, Atg8 (a master regulator of autophagosome elongation), or JNK, displayed resistance to bacterial infection (**Fig. 2C-D**). Importantly, trypan blue experiments demonstrated that cells in which target proteins had been knocked down displayed viability that was similar to controls treated with scrambled dsRNAs (Pandey et al., 2018; Qin et al., 2011; Qin et al., 2008). Thus, the differences in the numbers of recovered intracellular bacteria did not arise as a consequence of differences in the viability of host cells treated with dsRNAs. These data implicated the activities of several key proteins (including JNK) in controlling the subcellular trafficking, survival or replication of the pathogen in *Drosophila* S2 cells.

### Host factors identified in *Drosophila* S2 cell infection models function similarly in mammalian host cell infection models

We used mammalian alveolar macrophages or epithelial cells to validate findings from experiments using *Drosophila* S2 cells. First, results from Western blot experiments showed that JNK or p38 phosphorylation accompanied *A. baumannii* invasion of alveolar cells (**Fig. 3A**). Second, we showed that pre-treatment of lung epithelial TC-1 cells with compounds that inactivate JNK or PI3K signaling (SP60012 or wortmannin, respectively) reduced the internalization of *A. baumannii* into host cells (**Fig. 3B-C**). Pre-treatment of host cells with myriocin did not alter the invasion of the pathogen (**Fig. 3D**), as was observed in fly cells. However, depolymerization of host actin by pre-treating host cells with cytochalasin D did impair *A. baumannii* invasion of host cells (**Fig. 3D**). Importantly, trypan blue exclusion experiments revealed that the drug treatments did not induce host cell death during bacterial invasion, thereby suggesting that the reductions in bacterial internalization observed in the drug-treated cells resulted from the loss of the activities of the protein drug targets.

**Figure 3.**
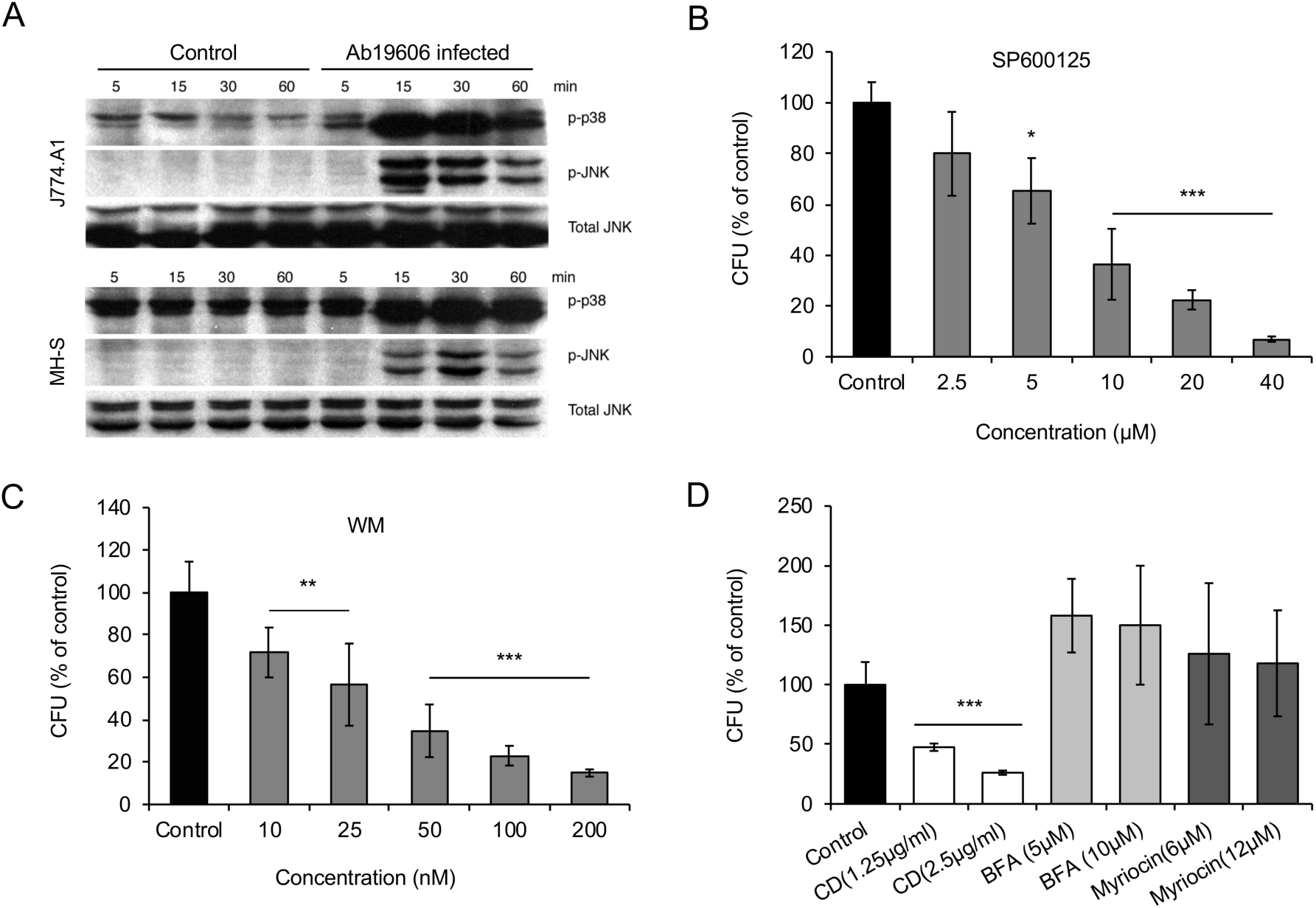
Host factors regulate *Acinetobacter baumannii* invasion of mammalian cells. (A) *A. baumannii* infection of the indicated macrophage cells for 1 hr (MOI = 50) stimulates host p38 and JNK phosphorylation. At the indicated time points, the infected host cells or uninfected controls were lysed for Western blotting analysis or CFU analysis. (B-D) Effects of pre-treating mouse lung epithelial TC-1 cells with pharmacological compounds SP600125 (B), wortmannin (WM) (C) or cytochalasin D (CD), brefeldin A (BFA) and myriocin (D) on Ab17978 invasion of host cells (1.5 h.p.i., MOI = 50). Data represent means ± SD from four independent experiments with triplicate wells examined for each treatment. *, **, ***: significance at p < 0.05, 0.01 and 0.001, respectively.

Finally, our fly cell findings indicated that proteins regulating organelle biogenesis and membrane trafficking contribute to controlling the intracellular lifestyle of the pathogen. To further explore this possibility, we used immunofluorescence microscopy to monitor the subcellular trafficking of *A. baumannii* in mammalian cells. We found that *A. baumannii* colocalized with EEA1 (early endosomal antigen 1)-containing membranes at early time points post-infection (≤ 1 h.p.i); interestingly, the compartment was also positive for the autophagosomal marker monodansylcadaverin (MDC) (Biederbick et al., 1995) (**Fig. 4A-C**), suggesting that the internalized bacteria are located in autophagic vacuoles. At later time points (4-24 h.p.i), a sub-population of the bacteria trafficked to a compartment that was decorated with the ER marker calreticulin (**Fig. 4D, G**). At the same time, a large sub-population trafficked to a compartment that contained lysosomal marker LAMP-1 or cathepsin D (**Fig. 4E-G**). These data indicated that a subpopulation of the pathogen avoided interactions with degradative lysosomal compartments and established a niche in an ER-like vacuole that were decorated with ER proteins. Taken together, findings from confocal immunofluorescence microscopy experiments performed with mammalian cells were consistent with insights garnered from experiments using the *Drosophila* S2 cell system.

**Figure 4.**
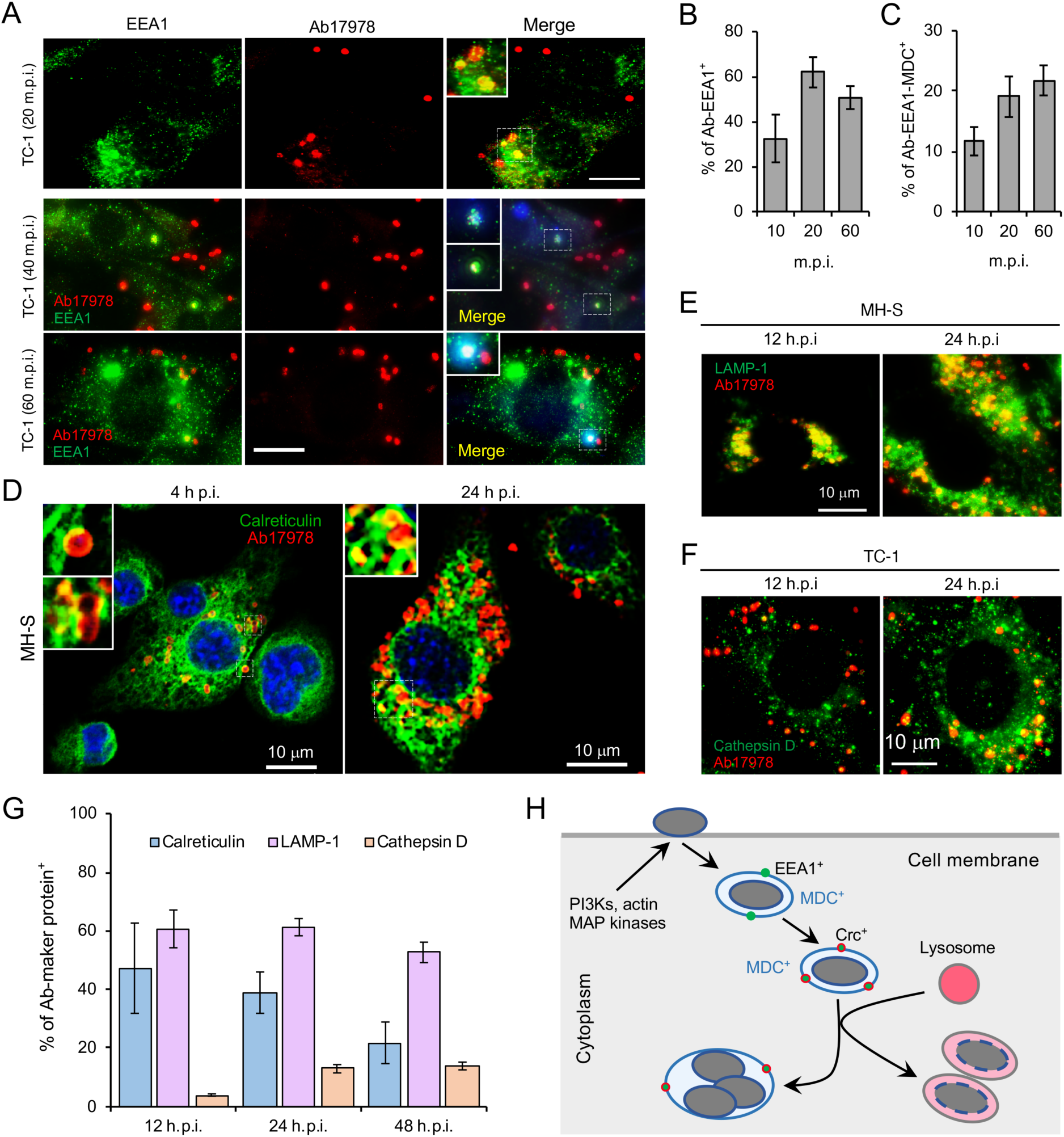
Intracellular trafficking and replication and/or persistence of *A. baumannii* in mammalian cells. Mouse macrophages (MH-S) and lung epithelial cells (TC-1) were infected with *A. baumannii* at an MOI of 20. At the indicated time points post infection, the cells were fixed with 3.7% formaldehyde and stained with the indicated antibodies and processed immunofluorescence microscopy assay. (A) Analysis of *A. baumannii* localization in host cells strained with the early endosomal marker EEA1 (early-endosomal autoantigen 1) and/or autophagosomal marker monodansylcadaverine (MDC) after internalization. (B, C) Quantification of *A. baumannii* colocalization with EEA1 (B) and with both EEA1 and MDC (C) at the indicated time points post-infection. (D) *A. baumannii* infected alveolar MH-S macrophages (D) were fixed and stained antibodies that recognize calreticulin, a host cell ER (endoplasmic reticulum) marker. (E, F) Localization of intracellular bacteria in host cells stained with antibodies that recognize the lysosome marker LAMP1 (E) or cathepsin D (F) at 12 and 24 h.p.i.. (G) Quantification of *A. baumannii* colocalization with calreticulin, LAMP1 or cathepsin D. (H) A proposal model of *A. baumannii* intracellular trafficking and persistence or replication. Bars: 10 μm. Representative confocal images showing the colocalization of *A. baumannii* (red) with the indicated host protein markers (green) inside MH-S macrophages. Quantitative data represent means ± SD from four independent experiments with triplicate wells examined for each treatment.

## Discussion

The genetically tractable *Drosophila S2* cell system possesses innate immune functions that are similar to mammalian cell systems. This attribute, together with efficient RNAi-mediated protein depletion and the availability of genome-scale RNAi libraries, has made the *Drosophila* S2 cell system a powerful platform for rapidly identifying host factors in large-scale screening campaigns (Qin et al., 2011; Qin et al., 2008). *Drosophila* S2 cells are also more resistant to *A. baumannii*-induced host cell death than their mammalian counterparts. The observed resistance facilitates analysis of host cell functions. Finally, the most important criterion for judging the utility of an alternative non-mammalian host-pathogen interaction model system is whether it can shed light on interactions that take place in cells from target organisms (e.g., mammalian cells). In this report, we demonstrate that host factors identified in the *Drosophila* cell system, including MAP kinases and proteins that regulate organelle biogenesis and membrane trafficking, contribute to regulating the internalization, intracellular trafficking and replication or persistence of bacterial pathogens in mammalian cells. Therefore, these findings not only demonstrate that the *Drosophila* S2 cell-*A. baumannii* interaction system recapitulates critical aspects of bacterial infection of mammalian cells, but also demonstrate the utility of the system.

*A. baumannii* is not predominately an intracellular pathogen. However, the bacterium can invade and persist in mammalian cells, including human lung, laryngeal, and cervical epithelial cells, and thereby promote invasive disease (Harding et al., 2018; Mortensen and Skaar, 2013). Following entry into host cells, the bacterium resides in a membrane-bound vacuole (Choi et al., 2008). Clathrin and β-arrestins are also engaged during the uptake into human lung epithelial cells (Smani et al., 2012). Similar to these findings, we found that upon internalization, *A. baumannii* resides in vacuoles decorated with the early endosome marker EEA1, or with both EEA1 and the autophagosomal marker MDC. During intracellular trafficking and persistence, some bacteria are observed in calreticulin-positive compartments. However, most intracellular bacteria locate in LAMP-1 or cathepsin D-positive vacuoles. Interestingly, intracellular persistence of *A. baumannii* was observed while the bacterium colocalized with lysosome, which is consistent with a previous report (Parra-Millan et al., 2018). The destabilization of lysosomal membranes (resulting in the release of cathepsin D outside of bacterium-containing lysosomes and the loss of acidic conditions inside lysosomes) may account for the intracellular persistence of the bacterium (Parra-Millan et al., 2018).

*A. baumannii* infection induces host cell autophagy, an evolutionary conserved self-eating process that plays critical roles in maintaining cellular homeostasis and host cell defense against microbial infections. Outer membrane protein A (OmpA) is a virulence factor of *A. baumannii* associated with the bacterial survival and pathogenicity (Sato et al., 2017). The bacterial proteins Omp33-36 can induce incomplete autophagy and assist intracellular replication of *A. baumannii* (Rumbo et al., 2014). Different pathways are activated during *A. baumannii* induced autophagy, including the AMPK/ERK/mTOR signaling pathway (Wang et al., 2016), the MAP/JNK signaling pathway (An et al., 2019) and the transcription factor EB (TFEB) pathway (Parra-Millan et al., 2018). Moreover, activation of TFEB could induce the autophagosome-lysosome system and destroys the acidic environment of the lysosome. These changes facilitate *A. baumannii* invasion, intracellular trafficking and persistence (Parra-Millan et al., 2018). We also found that infection increases the phosphorylation levels of host cell JNK and p38 (**Fig. S2A**), molecules that control (in part) autophagosome biogenesis (Sui et al., 2014). Similar to previous findings (Parra-Millan et al., 2018), both *Drosophila* S2 and mammalian host cells treated with bafilomycin, an inhibitor of organelle (e.g., endosomes and lysosomes) acidification and fusion between autophagosomes and lysosomes (Rumbo et al., 2014), wortmannin, or SP600125, reduced *A. baumannii* invasion. In the presence of bafilomycin, replication efficiency of *A. baummanii* increases, which is consistent with the observation of *A. baumannii* replication within autophagosomes (Rumbo et al., 2014). Taken together, we found that the intracellular lifestyle of *A. baumannii* is controlled by factors that regulate host cell autophagosome and autophagolysosome biogenesis (**Fig. 4H**). Importantly, the role of these proteins in supporting *A. baumannii* intracellular survival and replication found in both mammalian and *Drosophila* cells. These findings further support idea that the *Drosophila* cell system can serve as a useful model system for elucidating host-*A. baumannii* interactions.

The mechanisms of *A. baumannii* intracellular persistence are poorly understood. Host TFEB is required for bacterial invasion, intracellular trafficking and persistence inside human lung epithelial A549 cells. After infection by the bacterium, host TFEB is activated which results in activation of lysosomal biogenesis and autophagy that could promote the death of host cells. Therefore, TFEB is associated with host defense against bacterial infection. Moreover, HLH-30, the TFEB orthologue of *Caenorhabditis elegans*, plays an important role in promoting survival of the nematode during infection by the bacterium (Parra-Millan et al., 2018). These findings demonstrate that the functions of host factors are conserved in the pathogenesis of *A. baumannii* in phylogenetically divergent organisms. Our data also demonstrate that host factors that mediate *A. baumannii* invasion, intracellular trafficking and persistence are functionally conserved in *Drosophila* S2 and mammalian host cells. Therefore, the *Drosophila* S2 cell-*A. baumannii* interaction system may facilitate identifying and elucidating host factors that regulate the bacterium infection.

In summary, our findings establish a novel model system for defining new host factors that regulate *A. baumannii* invasion, intracellular trafficking and replication or persistence. By reporting host factor activities that, when disrupted, suppress intracellular survival of the pathogen, this work opens up new research avenues for defining potential therapeutics that target host cell functions. Therefore, this work will ultimately contribute to the fight against this harmful bacterial agent.

## Materials and Methods

### Cell Cultures and dsRNA-Mediated Knockdown of Target Genes

Mouse alveolar macrophage MH-S (ATCC CRL-2019), mouse lung epithelial cell TC-1 (ATCC, CRL-2785) and MLE12 (ATCC CRL-2110) were routinely cultured at 37°C in a 5% CO_2_ atmosphere in Dulbecco’s Modified Eagle’s Medium (DMEM) supplemented with 10% fetal bovine serum (FBS). *Drosophila* S2 cell cultivation and dsRNA-mediated knockdown of target genes in *Drosophila* S2 were performed as previously described (Pandey et al., 2018; Qin et al., 2008). Cells were seeded in 24-well plates and cultured overnight prior to infection.

For antibiotic protection assays, 2.5×10^5^ cells were seeded in each well; for fluorescence microscopy assays (see below), 1.0×10^5^ cells were seeded on 12-mm glass coverslips with a thickness of 0.13 mm (Fisherbrand^®^) placed on the bottom of 24-well plates.

### *Acinetobacter* Infection and Gentamicin Protection Assay

*Acinetobacter baumannii* wild-type strains ATCC17978 and ATCC19606 were used in this study. Bacteria were grown in LB broth or on LB agar plates. To prepare bacterial inoculums, 5 ml of LB broth was inoculated with a single colony from a freshly grown LB plate. Cultures were then grown at 37°C overnight with shaking. Bacteria were than washed with 1×PBS (pH 7.4) and re-suspended in PBS. Host cells were infected with *A. baumannii* at an MOI of 50. Briefly, infected cells were incubated at 37°C (28°C for *Drosophila* S2 cells, unless otherwise stated) after centrifugation for 5 min at 200×g. Thirty minutes post-infection, culture media were removed and infected cells were rinsed twice with PBS (pH7.4). Fresh media containing 50 μg/ml gentamicin were added to kill extracellular bacteria. Infected cells were continuously incubated in this antibiotic at 37°C. At various times post-infection, infected cells were lysed with 0.5% Tween 20 in sterile water, and the cell lysates were diluted in peptone saline [1% (wt/vol) Bacto peptone and 0.85% (w/v) NaCl]. Appropriate dilutions were spotted on LB plates and incubated at 37°C overnight to determine colony counts.

### Viability and Membrane Permeability Assays of Infected Host Cells

Cells cultured in 24-well plate were infected with *A. baumannii* as described above. The morphology of infected cells was observed and recorded using a Nikon (Eclipse Ti) laser confocal microscope. To quantitate cytotoxicity, cells cultured in 24-well plates were infected with *A. baumannii* in triplicate wells as described above. The culture supernatants were collected at various time points p.i., and the lactate dehydrogenase (LDH) release was determined by a CytoTox 96 nonradioactive cytotoxicity assay (Promega, Madison, Wis.) as described previously (Pei and Ficht, 2004; Pei et al., 2008). Cytopathic cell death is expressed as a percentage of maximum LDH release, i.e., 100 × (optical density at 490 nm [OD_490_] of infected cells - OD_490_ of uninfected cells)/(OD_490_ of lysed uninfected cells - OD_490_ of uninfected cells). A typan blue dye (0.2%) exclusion assay (Qin et al., 2008) was also employed to quantify the cytopathic effect or evaluate membrane permeability on mammalian and *Drosophila* S2 cells caused by *A. baumannii* infection or by drug or dsRNA treatment. Host cells in which dsRNA or drug treatment did not induce significant differences in viability and membrane permeability were used in subsequent experiments.

### Monodansylcadaverine (MDC) Staining

MDC staining was performed to evaluate the abundance of autophagic vacuoles in *A. baumannii* infected cells as previously described (Biederbick et al., 1995). Briefly, a 50 mM stock solution of MDC was prepared in dimethyl sulfoxide (DMSO). Cells were stained with MDC at a final concentration of 50 μM for 1 hr at room temperature, washed with phosphate-buffered saline for 3×10 min, and then examined by fluorescence microscopy.

### Immunofluorescence Microscopy

To determine where *A. baumannii* replication occurred, host cells were infected with *A. baumannii* and fixed with 3.7% formaldehyde fixed at 24 h.p.i.. The fixed cells were washed and processed for immunofluorescence microscopy (Pandey et al., 2017; Pandey et al., 2018; Qin et al., 2011; Qin et al., 2008). The primary antibodies used were as follows: mouse polyclonal anti-*Acinetobacter*, made by a commercial company, anti-EEA1, anti-LAMP1, anti-calreticulin, anti-cathepsin D (Santa Cruze, CA, USA). Samples were stained with Alexa Fluor 488-conjugated donkey anti goat IgG or Alexa Fluor 488-conjugated chicken anti-rabbit IgG and Alexa Fluor 594-conjugated donkey anti mouse (Molecular Probes). Cover slips were then mounted in Vectashield^®^ mounting media (Vector Laboratories, Inc., CA, USA) and visualized with the Nikon confocal microscope. Image acquisition and processing were performed as previously described (Pandey et al., 2017; Pandey et al., 2018; Qin et al., 2011; Qin et al., 2008).

### Drug Treatments

*Drosophila* S2 and/or murine cells were coincubated with assorted pharmacological compounds, including cytochalasin D (CD, actin polymerization inhibitor), wortmannin [WM, phosphoinositide 3-kinase (PI3K) inhibitor)], bafilomycin A (BAF), brefeldin A (BFA) Myriocin, SB203580 (MAP kinase p38 inhibitor) and SP600125 1 hr before, and during, infection with the indicated *A. baumannii* strains. Uptake of the microorganisms by the treated and untreated cells and intracellular replication and/or persistence of *A. baumannii* population were determined using the gentamicin protection assay as described above.

### Western Blot

To determine MAP kinase activation, cells cultured in 24-well plates were infected with *A. baumannii* at an MOI of 50. Uninfected cells were used as control. At 5, 15, 30 and 60 min post infection (m.p.i.), the infected and uninfected control cells were lysed and subjected to Western blotting analysis as previously described (Pandey et al., 2017; Pandey et al., 2018; Qin et al., 2011). Primary antibodies used in the analysis were: anti-p38, anti-SAPK/JNK, anti-p-SAPK/JNK (Cell signaling). Dilution of primary antibodies and secondary antibody (HRP anti-IgG, Sigma-Aldrich, USA) was 1:1,000 and 1:1,000∼5,000, respectively.

### Statistical Analysis

All quantitative data were derived from results obtained from at least three independent experiments with triplicate treatments tested. The data of controls were normalized as 1 or 100% to easily compare results from different independent experiments. The significance of the data was assessed using the Student’s t-test.

## Acknowledgments

PdF is supported by funding from the Texas AM Clinical Science Translational Research Institute Pilot Grant CSTR2016-1, DARPA (HR001118A0025-FoF-FP-006), NIH (R21AI139738-01A1, 1 R01AI141607-01A1, 1R21GM132705-01), the National Science Foundation (DBI 1532188, NSF0854684) and the Bill Melinda Gates Foundation. The project depicted was in part sponsored by the Defense Advanced Research Projects Agency (Agreement HR001118A0025-FoF-FP-006). This research was also supported by NIH grant awards NIH 1R01 AI48496-01A1 and NIH 1U54AI057156-0100 to TAF; grant awards from the National Natural Science Foundation of China (# 81371773) to QMQ. The content of the information does not necessarily reflect the position or the policy of the Government, and no official endorsement should be inferred.

## Author Contributions

QMQ, JP, ARF, TAF and PdF conceived and designed the experiments. QMQ, JP, and GG performed the experiments. QMQ, JP, PdF and TAF analyzed the data. PdF, ARF and TAF contributed reagents, materials, and analysis tools. QMQ and PdF wrote the paper.

