## Supplemental Figure S1 to S4 for "A tractable *Drosophila* cell system enables rapid identification of *Acinetobacter baumannii* host factors" (from another version of this preprint: version 1, 2020-04-10)

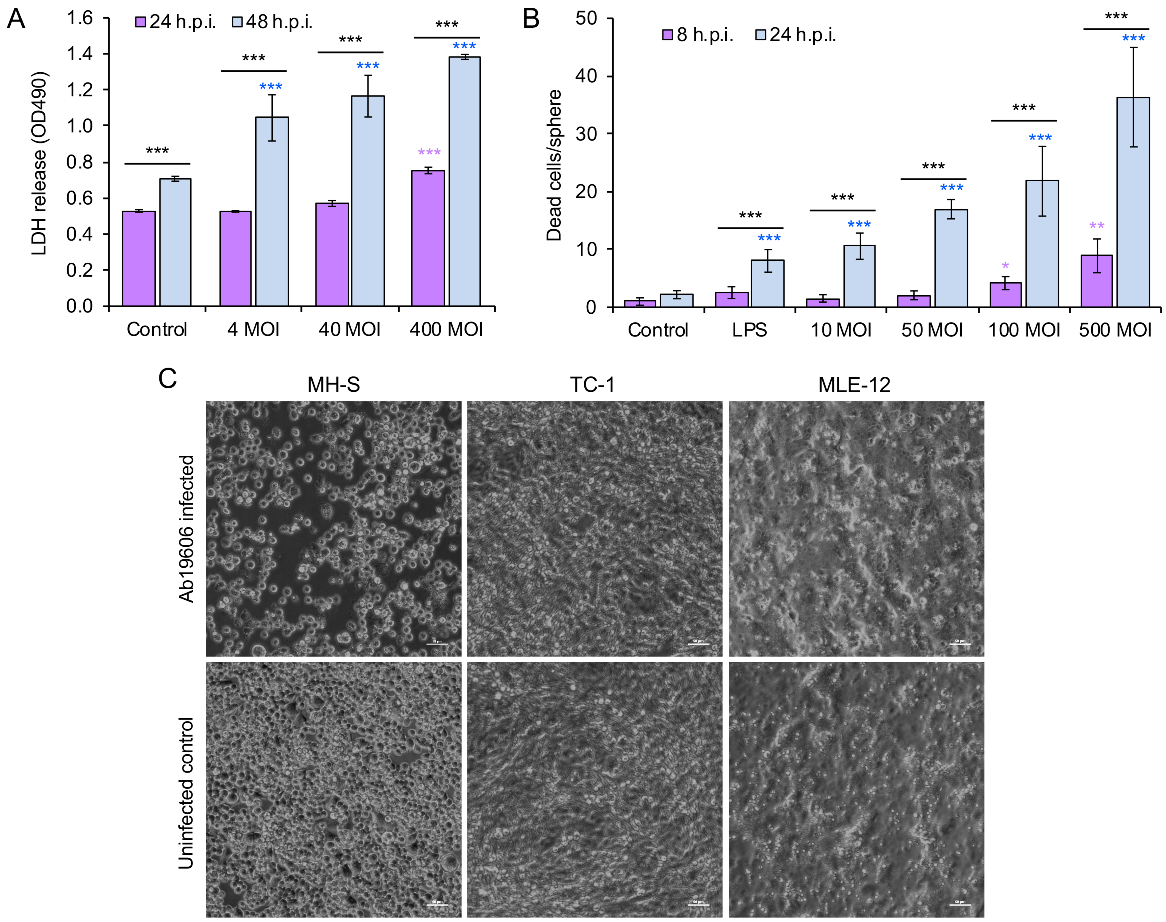


**Figure S1.** ***Acinetobacter baumannii*** **induces alveolar macrophage cell death.** (A, B) *A. baumanni* (Ab19609) infects lung alveolar macrophages and host cell dead is determined by LDH lease (A) or by microscopy measurement (B). Purple and blue asterisks: compared to the controls at the same time point post infection (the same color), respectively. h.p.i.: hr post infection; Black asterisks: compared to the same MOI at different time points. *, **, and ***: significance at $p < 0.05$, 0.01, and 0.001, respectively. Data represent means ± standard deviation (SD) from three independent experiments with triplicate wells examined for each treatment. (C) Morphology of *Acinetobacter*-infected and uninfected cells at 48 h.p.i.. Lung alveolar macrophages MH-S, epithelial cells TC-1 and MLE-12 cultured in 24-well tissue culture plates were infected with *A. baumannii* at an MOI (multiplicity of infection) of 50. The cells were observed by phase-contrast microscopy. Uninfected cells were used as control. Scale bar: 50 mm. Images shown are a representative of three independent experiments.

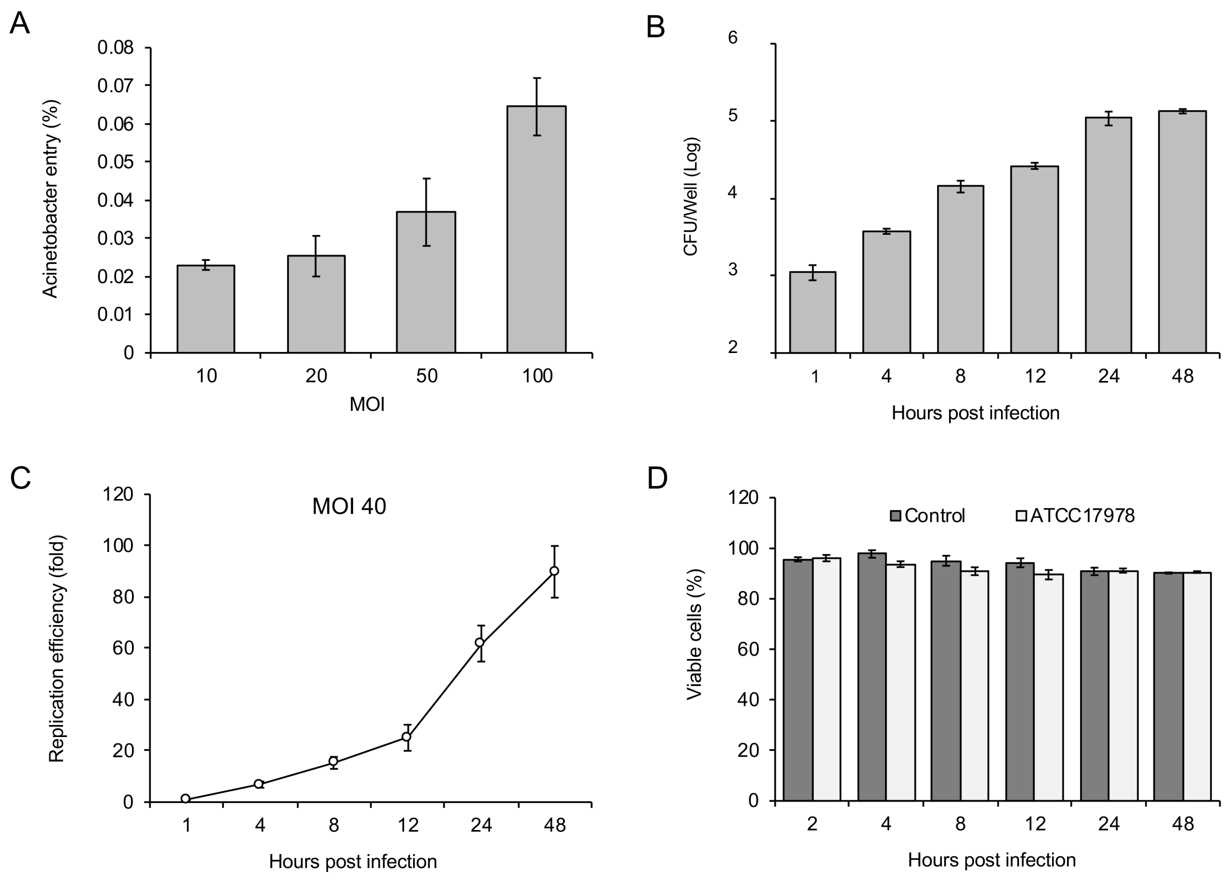


**Figure S2.** ***Acinetobacter baumannii*** **(ATCC 17978) invades and replicates in** ***Drosophila melanogaster*** **S2 cells.** (A) A. *baumannii* invasion of S2 cells was measured at the indicated MOIs. (B) Intracellular replication of Ab17978 in S2 cells during a time course (48 hr) of infection. (C) Intracellular replication efficiency of *A. baumannii* in *Drosophila* S2 cells. (D) Viablity assay of S2 cells infected by A. *baumannii* strain 17978. Data represent means ± SD from three independent experiments with triplicate wells examined for each treatment.

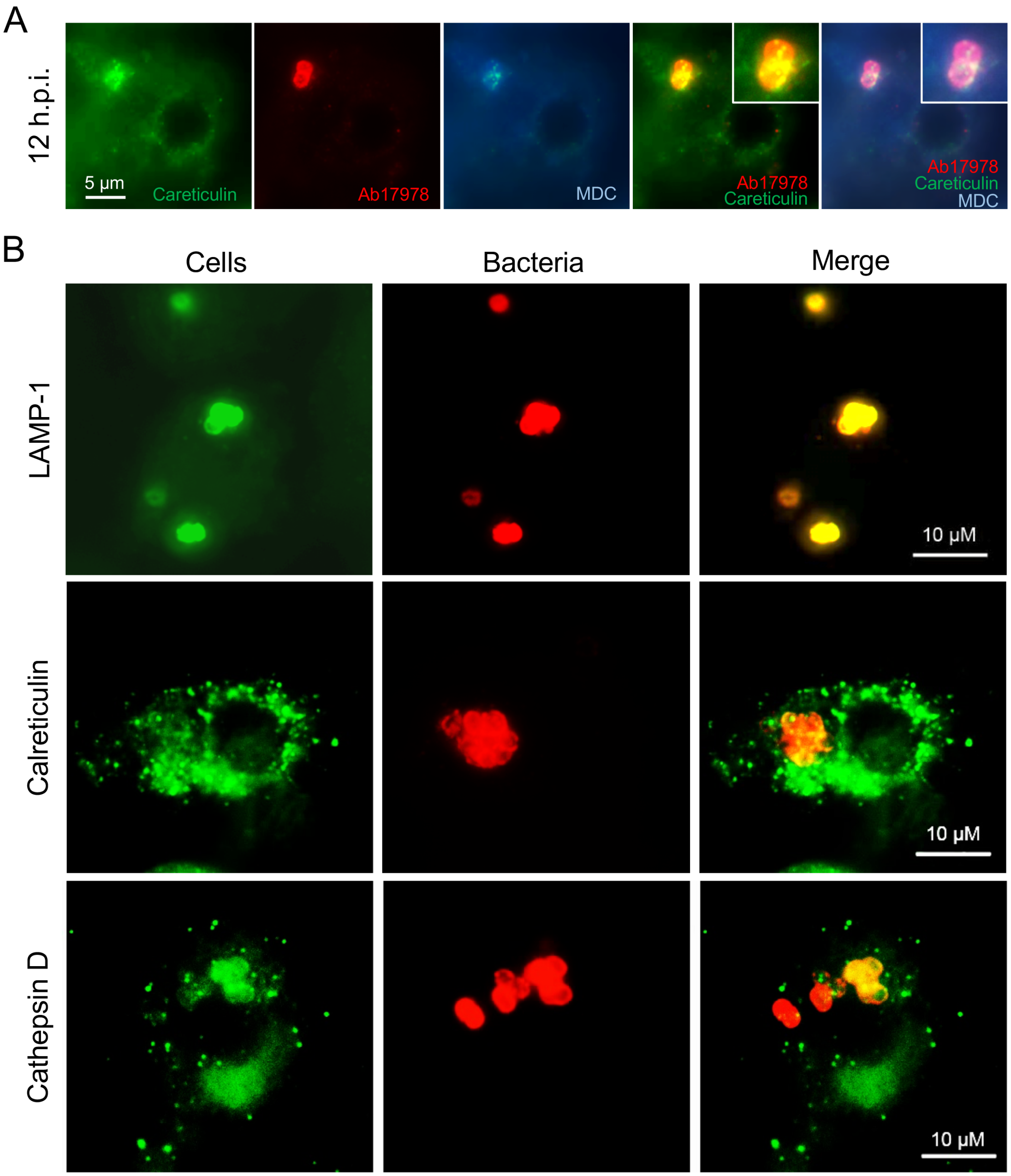


**Figure S3. Intracellular trafficking of *A. baumannii* (Ab17978) in *Drosophila* S2 cells.** (A) *Drosophila* S2 cells were infected with *A. baumannii* Ab17978 with an MOI of 20, and at 12 h.p.i., the infected cells were fixed and stained with ER (endoplasmic reticulum) marker calreticulin and with autophagosomal marker monodansylcadaverine (MDC). Insets: magnified the intracellular bacteria. (B) *A. baumannii* Ab17978 infected *Drosophila* S2 cells (24 h.p.i.) were stained with antibodies that recognize the lysosome marker LAMP1, cathepsin D, or ER marker calreticulin. Images are from a representative of three independent experiments.

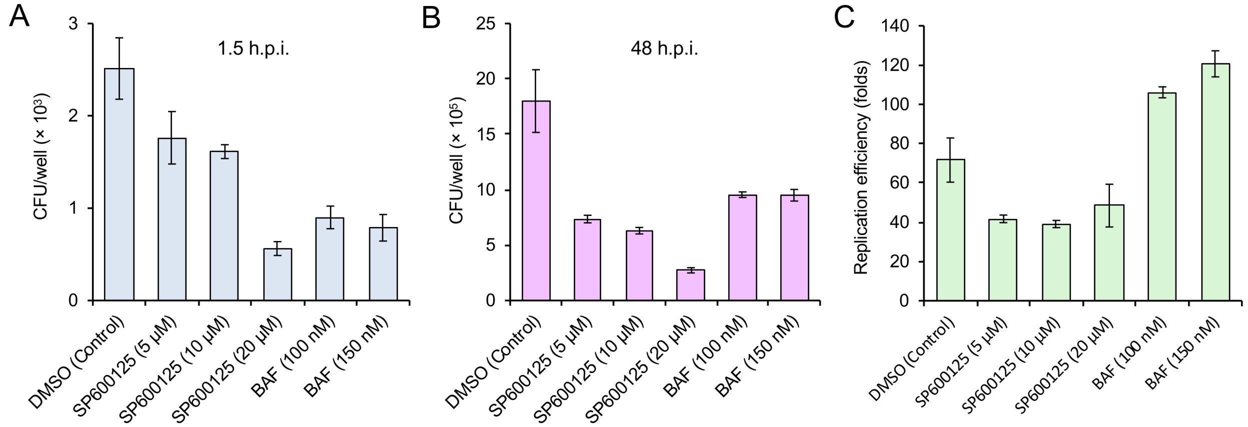


**Figure S4. Host cells treated with pharmacological compounds related to autophagy affect *A. baumannii* invasion and intracellular replication.** *Drosophila S2* cells treated with the indicated compounds before 1 hr and during the bacterial infection, at 1.5 and 48 h.p.i., the infected cells were lysed and subjected to gentamicin protection assays. (A, B) *A. baumannii* invasion (A) and intracellular replication (B) in a time course of 48 hr infection. (C) *Drosophila* S2 cells treated by the indicated compounds affect intracellular replication efficiency of *A. baumannii*. Data represent means ± SD from a representative experiment with triplicate wells examined.
